# A dynamic regulatory interface on SARS-CoV-2 RNA polymerase

**DOI:** 10.1101/2020.07.30.229187

**Authors:** Wei Shi, Ming Chen, Yang Yang, Wei Zhou, Shiyun Chen, Yi Yang, Yangbo Hu, Bin Liu

## Abstract

The RNA-dependent RNA polymerase (RdRp) of SARS-CoV-2 is the core machinery responsible for the viral genome replication and transcription and also a major antiviral target. Here we report the cryo-electron microscopy structure of a post-translocated SARS-CoV-2 RdRp core complex, comprising one nsp12, one separate nsp8(I) monomer, one nsp7-nsp8(II) subcomplex and a replicating RNA substrate. Compared with the recently reported SARS-CoV-2 RdRp complexes, the nsp8(I)/nsp7 interface in this RdRp complex shifts away from the nsp12 polymerase. Further functional characterizations suggest that specific interactions between the nsp8(I) and nsp7, together with the rearrangement of nsp8(I)/nsp7 interface, ensure the efficient and processive RNA synthesis by the RdRp complex. Our findings provide a mechanistic insight into how nsp7 and nsp8 cofactors regulate the polymerase activity of nsp12 and suggest a potential new intervention interface, in addition to the canonical polymerase active center, in RdRp for antiviral design.

**Author summary:** Since it was first discovered and reported in late 2019, the coronavirus disease 2019 (COVID-19) pandemic caused by highly contagious SARS-CoV-2 virus is wreaking havoc around the world. Currently, no highly effective and specific antiviral drug is available for clinical treatment. Therefore, the threat of COVID-19 transmission necessitates the discovery of more effective antiviral strategies. Viral RNA-dependent RNA polymerase (RdRp) is an important antiviral drug target. Here, our cryo-EM structure of a SARS-CoV-2 RdRp/RNA replicating complex reveals a previously uncharacterized overall shift of the cofactor nsp8(I)/nsp7 interface, leading to its rearrangement. Through *in vitro* functional test, we found that the specific interactions on the interface are important to the efficient RNA polymerase activity of SARS-CoV-2 RdRp. These observations let us to suggest this interface as a potential new drug intervention site, outside of the canonical polymerase active center, in RdRp for antiviral design. Our findings would provide new insights into regulatory mechanism of this novel SARS-CoV-2 RdRp, contribute to the design of antiviral drugs against SARS-CoV-2, and benefit the global public health.

## INTRODUCTION

The novel severe acute respiratory syndrome coronavirus-2 (SARS-CoV-2) is the causative agent of the coronavirus disease 2019 (COVID-19) and has resulted in a worldwide pandemic with over 15 million infections [1-4]. SARS-CoV-2 is an enveloped, positive-sense single-stranded RNA virus. As a new member of Betacoronavirus genus, SARS-CoV-2 appears to be more transmissible than other members [5-7]. The threat of COVID-19 pandemic demands a prompt effective antiviral strategy.

The RNA-dependent RNA polymerase (RdRp) of SARS-CoV-2 is the key machinery responsible for the viral RNA synthesis, and thus has become a promising target for nucleoside analogue inhibitors, such as Remdesivir and EIDD-2801 [8-10]. The RdRp complex of SARS-CoV-2 comprises the core catalytic non-structural protein (nsp) 12, and the cofactors nsp7 and nsp8 that reinforce the RNA template binding and processive RNA synthesis [11, 12]. Recent cryo-electron microscopy (cryo-EM) structures of the SARS-CoV-2 apo RdRp and RdRp/RNA complexes [13-17] have revealed that their overall architectures are highly similar to that of SARS-CoV RdRp [11]. Here we present the cryo-EM structure of SARS-CoV-2 RdRp complex in post-translocation state during viral RNA synthesis at near-atomic resolution. The structure reveals a previously uncharacterized overall rearrangement of nsp8(I)/nsp7 interface, which is critical to RNA synthesis in *in vitro* transcription assays, and suggests a potential new antiviral intervention site.

## RESULTS

### Overall architecture of RdRp/RNA complex

To obtain the target structure, we incubated the individually purified nsp12, nsp7 and nsp8 proteins in the presence of a template-primer RNA duplex and adenosine-5′-triphosphate (ATP) to allow one nucleotide to be incorporated into the 3′ end of the primer chain. The resulting RdRp/RNA complex was used in cryo-EM imaging and structural determination. More than 4 million initial particles were automatically picked from 4,750 micrograph movies and 123,246 particle projections were used to reconstruct the final overall 3.4 Å resolution density map of the complex (S1 Fig and S1 Table).

The structure of RdRp/RNA complex contains one nsp12, one nsp7 and two nsp8 subunits, nsp8(I) and nsp8(II), and displays similar overall configuration to that observed in the recently reported structures (Fig 1A-1C) [13, 15]. The cryo-EM map contains unambiguous density for most parts of nsp12, nsp7 and nsp8 except for the N-terminal 88 residues of nsp12, N-terminal 77 residues of nsp8(I), C-terminal 20 residues of nsp7, and 167 residues of nsp8(II), which are not modeled in the cryo-EM map due to poor density (Fig 1A-1C and S2A-S2C Fig). The nsp12 protein comprises an N-terminal β-hairpin (residues 29-50) [13] and a nidovirus RdRp-associated nucleotidyltransferase domain (NiRAN, residues 84-249) [18], which is connected by an interface domain (residues 250-365) to the polymerase domain (residues 366-918). Like RdRps from other RNA viruses, nsp12 polymerase domain adopts an encircled right hand architecture with fingers (residues 366-581 and 621-679), palm (residues 582-620 and 680-815) and thumb (residues 816-918) subdomains surrounding the active site (Fig 1B and 1C) [19, 20]. The encircled polymerase domains are further stabilized by nsp8(I) subunit and nsp7-nsp8(II) subcomplex. The nsp8(I) sits upon the fingers and interface domains, and nsp7-nsp8(II) subcomplex presses part of the index finger (residues 408-446 of nsp12) against the thumb domain (Fig 1C). The template-primer RNA duplex is held in the RNA exit channel by an extended loop from the fingers domain and an α-helix from the thumb domain (Fig 1C). This RNA-binding mode is also adopted by other positive-sense RNA virus RdRps [21-23].

**Fig 1.**
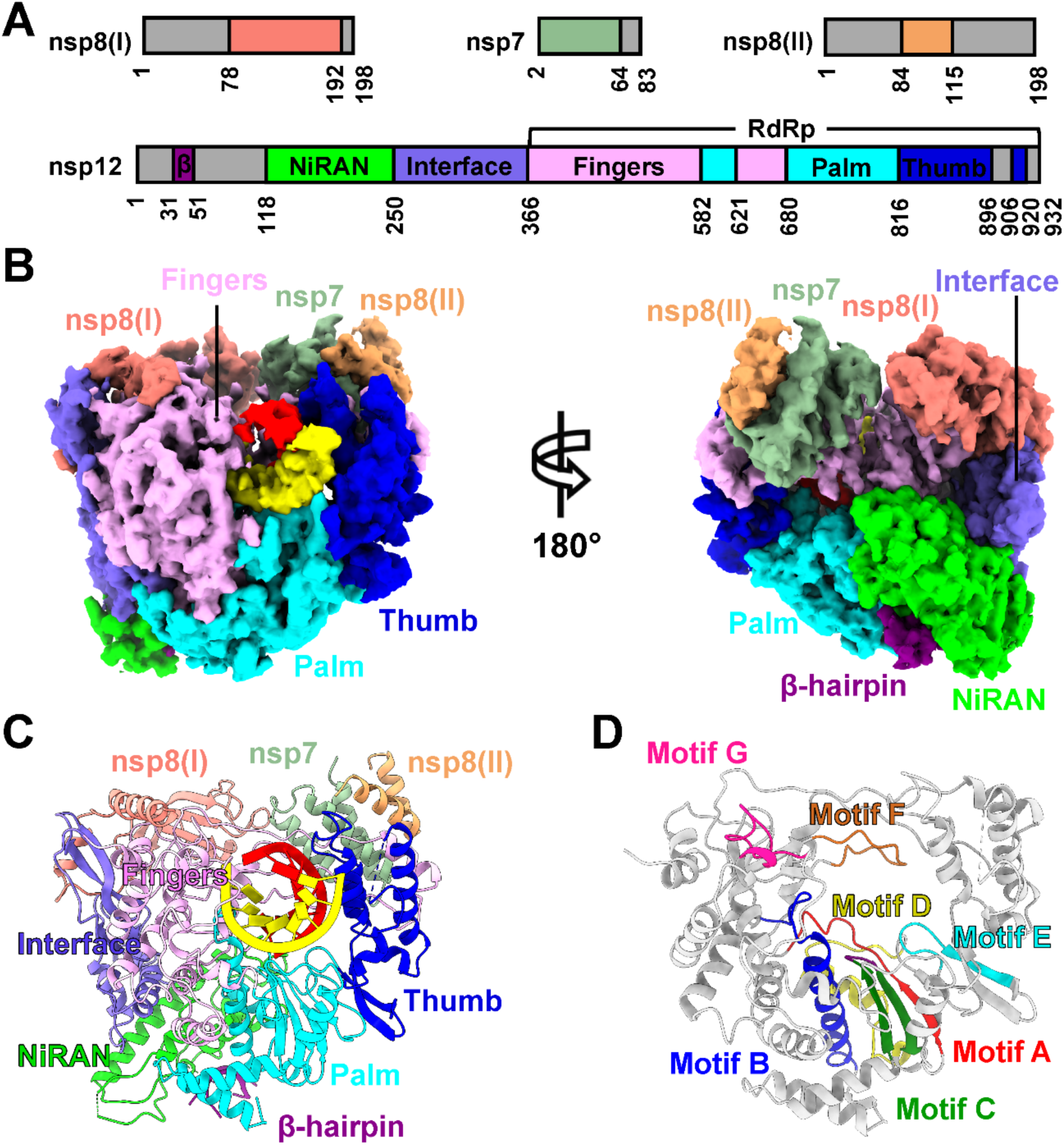
Cryo-EM structure of SARS-CoV-2 RdRp/RNA complex. **(A)** Schematic diagram of SARS-CoV-2 nsp7, nsp8 and nsp12 proteins in RdRp complex. The portions that are not modeled in the map are colored in gray. **(B** and **C)** Two directional views of the split cryo-EM map (**B**) and the model (**C**) of RdRp/RNA complex, contoured at 0.1 of view value of Chimera. The color scheme is consistent with Fig 1A. **(D)** Spatial arrangement of seven motifs A-G surrounding the active site. Only nsp12 polymerase domain is shown, and other parts of the complex are omitted for clear presentation.

### A post-translocated catalytic active site

The catalytic core of nsp12 polymerase domain is formed by motifs A-G (Fig 1D and S2D-S2F Fig), which have the conserved spatial arrangement for all viral RdRps [24, 25]. Residue D618 of motif A and residues D760 and D761 of motif C are involved in the coordination of catalytic magnesium ions. The observed active site in this RdRp/RNA complex is in a post-translocation state, in which the polymerase active site contains one magnesium ion that is coordinated by carboxyl groups of residues D618 and D761 (Fig 2A). The magnesium ion was not built in the recent post-translocated SARS-Cov-2 RdRp/RNA complex [17] (Fig 2B). The position of Mg^2+^ in this study is spatially similar to that in Enterovirus 71 (EV71) RdRp/RNA post-translocated elongation complex (Fig 2C) [22]. By contrast, two catalytic magnesium ions are coordinated by the three active site aspartate residues in the pre-translocated SARS-CoV-2 RdRp/RNA/Remdesivir complex [15] (Fig 2D) and the Hepatitis C virus (HCV) RdRp/RNA/Sofosbuvir complex (Fig 2E) [26], whereas in the apo RdRp of SARS-CoV-2 [13], no divalent cation was identified in the active site (Fig 2F) and three aspartate residues adopt different rotamer conformations compared to those in the RNA-bound complexes. These structures suggest an evolutionarily conserved nucleotide addition mechanism by utilizing two divalent cations for all polymerases, which are important for drug design targeting viral RdRps [27].

**Fig 2.**
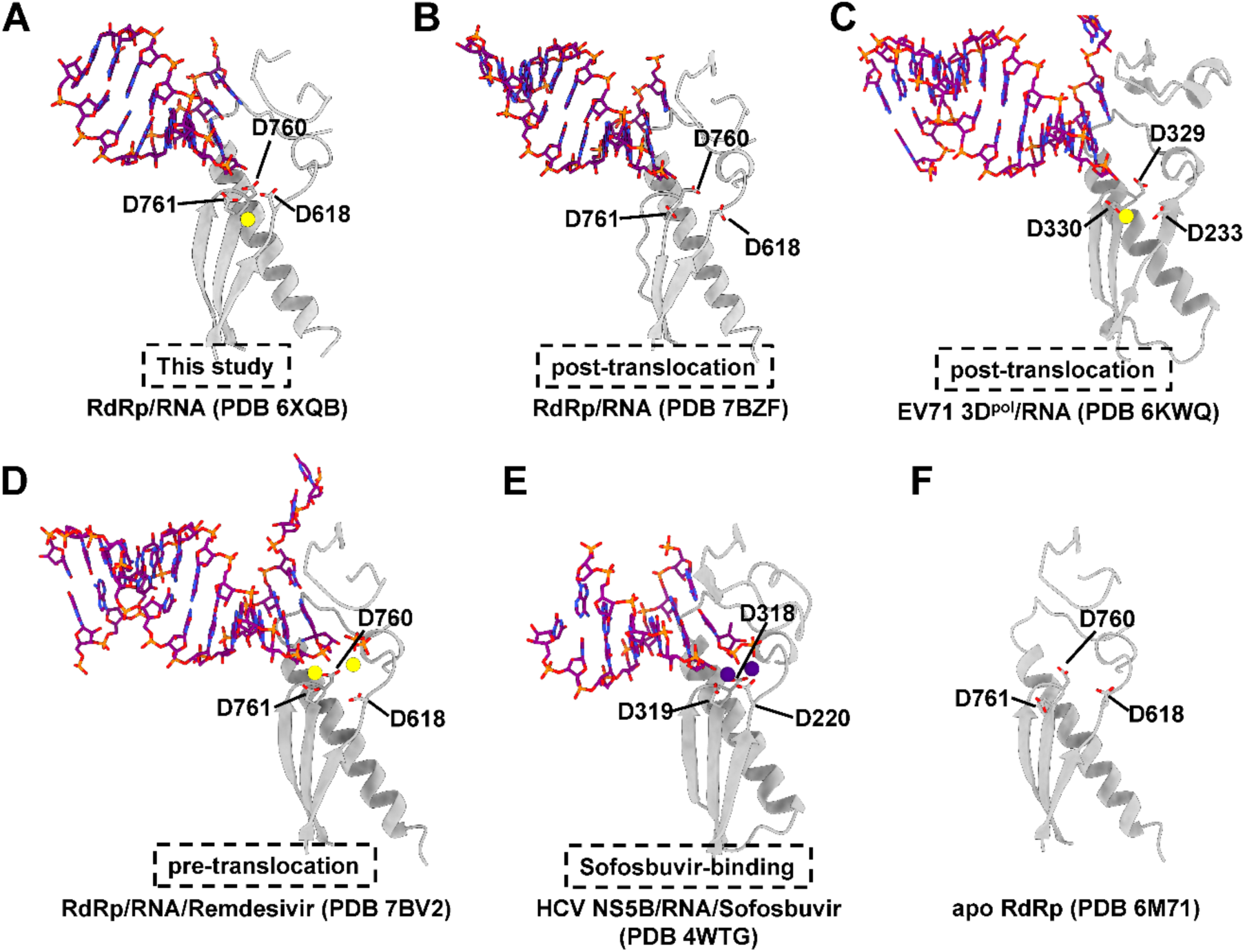
Structural comparison of catalytic active sites among multiple RdRp complexes. **(A-F)** Detailed interactions between three conserved aspartic acids at motifs A and C and the coordinated divalent metal ions in this SARS-CoV-2 post-translocated RdRp/RNA complex (**A**), other SARS-CoV-2 post-translocated RdRp/RNA complex (**B**), EV71 post-translocated 3D^pol^/RNA complex (PDB 6KWQ, **C**), SARS-CoV-2 pre-translocated RdRp/RNA/Remdesivir complex (PDB 7BV2, **D**), HCV Sofosbuvir-binding NS5B/RNA complex (PDB 4WYG, **E**) and SARS-CoV-2 apo RdRp (PDB 6M71, **F**). The conserved aspartic acids involved in the catalysis are shown as sticks. Magnesium and manganese ions are show as yellow and purple balls, respectively.

### Structural comparisons reveal rearrangement of nsp8(I)/nsp7 interface

Structural comparison between this RdRp/RNA complex and SARS-CoV-2 apo RdRp (PDB 6M71) [13] reveals a relatively stable overall conformation of the polymerase throughout the nucleotide addition cycle (Fig 3A), which was also observed in other viral RdRps [26, 28]. One evident difference in the polymerase structures between the RNA-bound form and the apo form is the outward movement of the thumb subdomain. In particular, the outmost α-helix shifts by ∼2.4 Å, slightly widening the RNA exit channel to accommodate the RNA duplex (Fig 3A). Notably, a ∼3.1 Å movement of the nsp7-nsp8(II) subcomplex away from RdRp polymerase domain and an accompanying ∼2.5 Å outward movement of the nsp8(I) C-terminal portion on the interface (Fig 3B) in this post-translocated complex lead to a rearranged interface between nsp8(I) and nsp7 (Fig 3C). The movements of both nsp8(I) and nsp7-nsp8(II) subcomplex in the RdRp complex are also noticeable when compared to the pre-translocated RdRp/RNA/Remdesivir complex (PDB 7BV2) [15] (Fig 3D). Structural comparisons of the nsp8(I)/nsp7 interfaces among the different SARS-CoV-2 RdRp/RNA complexes [13, 15-17] by superimposition on the nsp7 also reveal shifts of nsp8(I) relative to nsp7 by ∼1-2 Å (S3A-S3D Fig), suggesting the slight opening of the interface in this structure.

**Fig 3.**
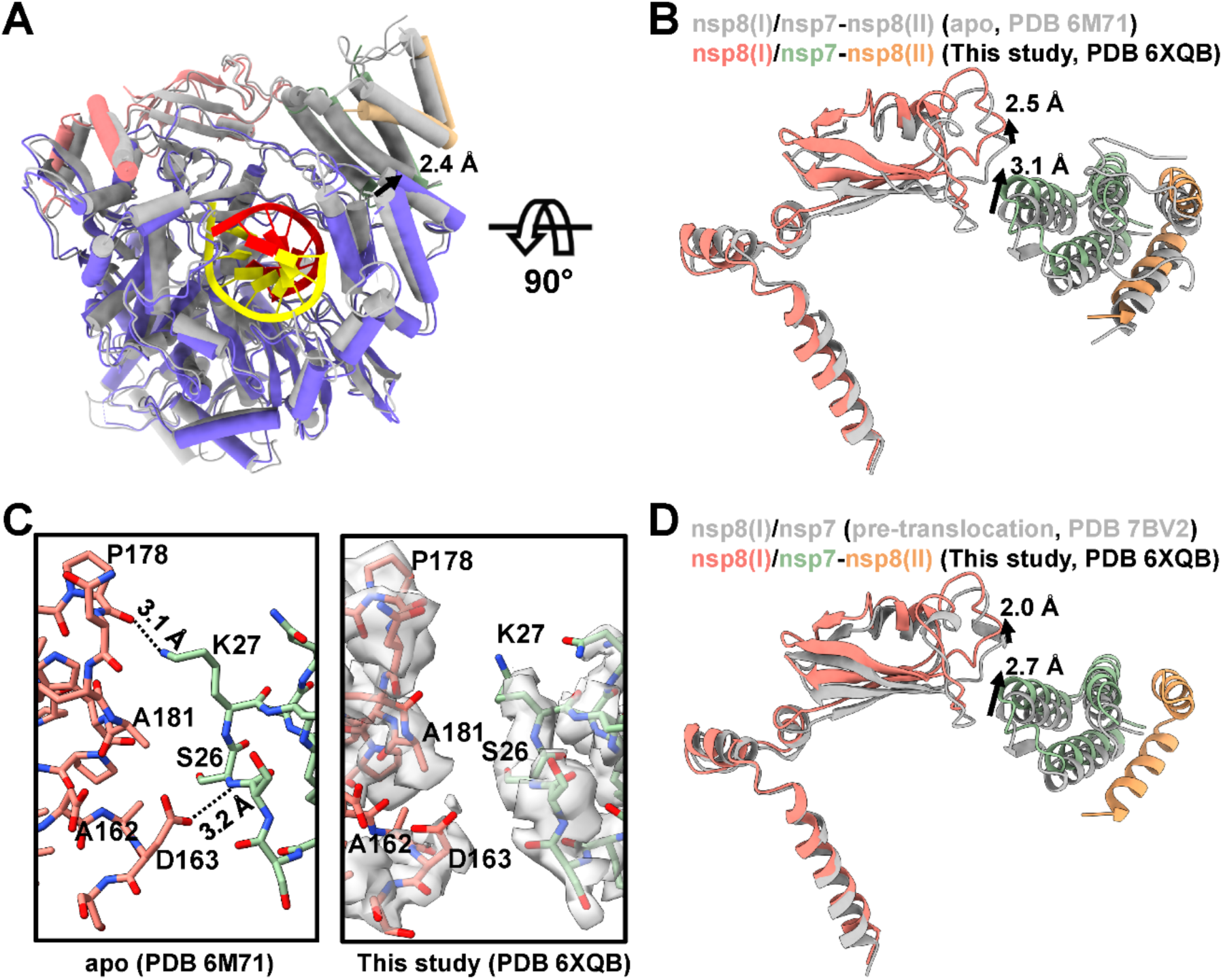
Structural differences at the interface between nsp8(I) and nsp7. **(A)** Superimposition of this RdRp/RNA complex with the apo RdRp (PDB 6M71, gray) via nsp12. **(B)** Closed-up view of the superimposed nsp7 and nsp8 cofactors. The distance shifts are indicated by black arrows and corresponding values. **(C)** Closed-up views of the interface between nsp8(I) and nsp7 in apo RdRp (left) and this post-translocated RdRp/RNA complex (right) with transparent split cryo-EM map, contoured at 0.1 of view value of Chimera. The nsp8(I) and nsp7 are colored in salmon and dark sea green, respectively. Residues that contribute to the interface interactions, as well as two neighboring residues are labelled. **(D)** Closed-up view of the nsp7 and nsp8 cofactors of this RdRp/RNA complex and the pre-translocated RdRp/RNA complex (PDB 7BV2, gray) by superimposition on nsp12.

### The nsp8(I)/nsp7 interface plays a critical role in RNA synthesis

In the apo RdRp structure (PDB 6M71) [13], residue D163 of nsp8(I) is hydrogen bonded with the main-chain nitrogen atom of S26 in nsp7, and K27 of nsp7 forms a hydrogen bond with the main-chain carbonyl group of P178 in nsp8(I). However, these interactions are abolished by the structural rearrangements of both nsp8(I) subunit and nsp7-nsp8(II) subcomplex in the post-translocated complex (Fig 3C). Further sequence alignment showed that these two residues D163 (nsp8(I)) and K27 (nsp7) are highly conserved in coronaviruses from all four genera (S4 Fig). Alanine substitutions of either D163 (nsp8(I)) or K27 (nsp7), which are supposed to weaken the interactions between nsp8(I) and nsp7 (Fig 4A) significantly decreased their abilities to activate the polymerase activity of nsp12 on a 5′ U_10_ RNA hybrid [15] (Fig 4B and 4C). In addition, replacing the alanine residue at 162 position by lysine (A162K) or at 181 position by asparagine (A181N) in nsp8, both of which are supposed to strengthen the nsp8(I)/nsp7 interaction through hydrogen bonds with S26 of nsp7 (Figs 3C and 4A), also showed decreased RNA extension activity (Fig 4D), suggesting an intricate regulation of nsp8(I)/nsp7 interaction to nsp12 activity. Addition of nsp7 and nsp8 proteins strongly increased the binding of RdRp complex to RNA hybrid [12] (S5A Fig), and none of the mutated nsp7 and nsp8 cofactors significantly influenced the RNA-binding affinity, as similar concentration of RdRp wild-type or mutated complex was required for shifting RNA (Fig 4E and 4F). Similarly, the decreases in RNA extension activities caused by interfering nsp8(I)/nsp7 interactions were also observed in the analyses based on a short RNA substrate (S5B and S5C Fig). Summarily, these results suggest that the nsp8(I)/nsp7 interface plays a critical role in stabilizing the polymerase domain and ensuring the processive RNA synthesis.

**Fig 4.**
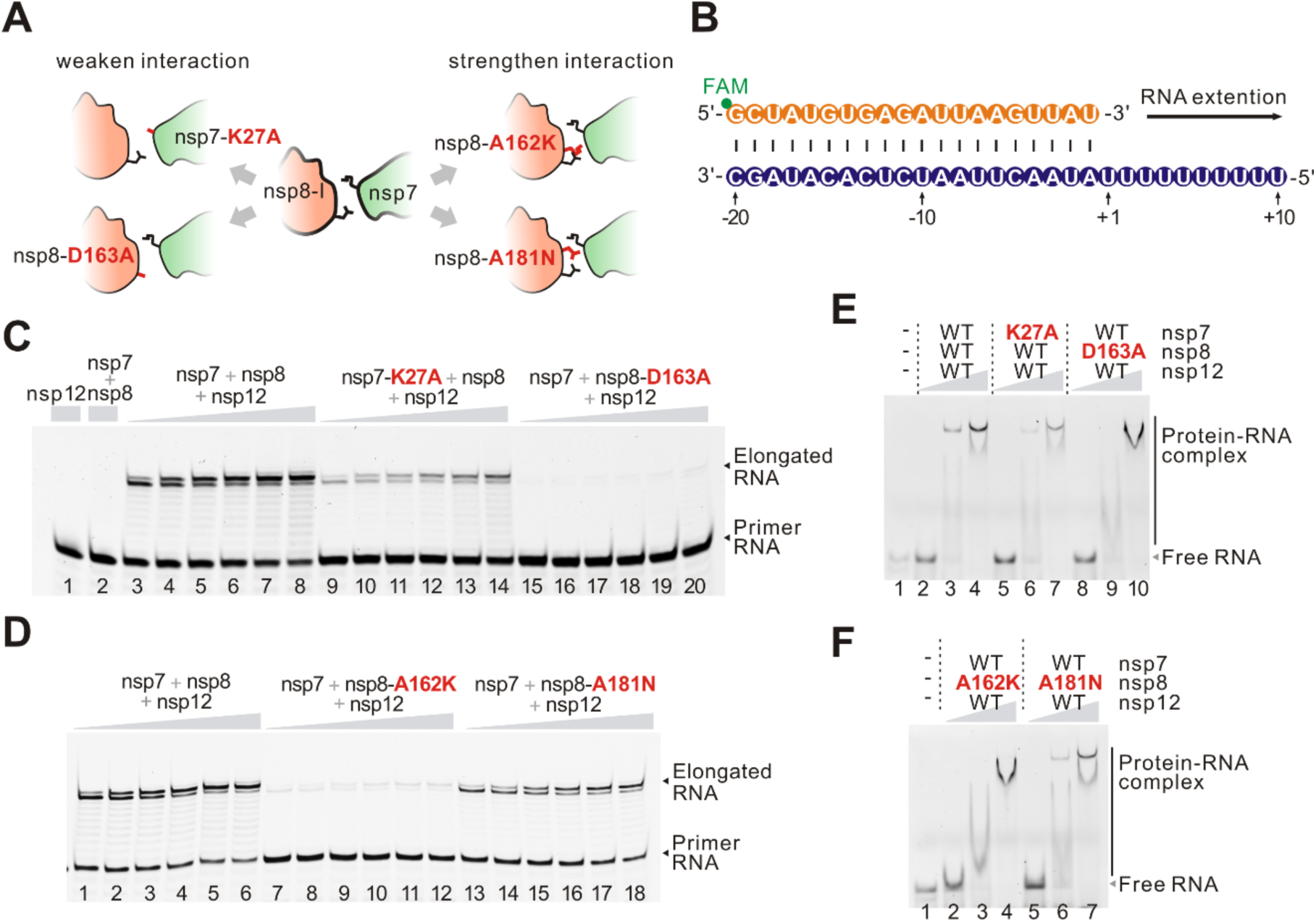
Role of the interface between nsp8(I) and nsp7 in RdRp polymerase activity. **(A)** Model of mutations introduced in nsp8 and nsp7 proteins to weaken or strengthen the nsp8(I)/nsp7 interaction. Mutated residues in the model are indicated in red. **(B)** Sequence of 5′ U_10_ RNA duplex used for RNA extension and RdRp/RNA complex formation tests. **(C** and **D)** Activity of recombinant RdRp complexes in RNA extension. Reactions were performed for 15s, 30s, 1 min, 2 min, 5 min and 10 min at 30 °C. Reactions for nsp12 alone or nsp7-nsp8 complex were carried out for 10 min. RdRp complexes with weakened nsp8(I)/nsp7 interaction are shown in panel **C** and those with strengthened nsp8(I)/nsp7 interaction are shown in panel **D. (E** and **F)** RNA binding of RdRp complexes with decreased (**E**) or increased (**F**) nsp8(I)/nsp7 interaction. RdRp complexes were used at concentrations of 200 nM, 400 mM and 800 nM. Each test was independently repeated more than three times with similar results.

## DISCUSSION

Since the short RNA duplex used in this study does not make direct contacts with cofactors nsp8 or nsp7 in the complex, the observed rearrangement in nsp8(I)/nsp7 interface is likely triggered by the minor conformational change of thumb domain during the polymerization reaction (Fig 3A). In addition, structural comparison of several reported nsp7-nsp8 heterodimers [29, 30] demonstrates that the N-terminal extension of nsp8 is highly flexible (S6 Fig), which may explain its invisibility in most coronavirus RdRp structures [11, 13, 15]. The invisible N-terminal extensions of nsp8 in this complex might contact the polymerase domain and thus allosterically transmit the motion in the polymerase domain to the nsp8(I)/nsp7 interface. Two recent studies presented structures of SARS-CoV-2 RdRp/RNA replicating complex in which the nsp8 helical extensions stabilize the exiting RNA and act as a ‘sliding poles’ to enable processive elongation of the long genome [16, 17]. As RNA elongates and exits from the channel, the stabilizing ‘sliding poles’ likely undergo some periodic movements during the processive RNA synthesis. Such movements could be transmitted to the nsp8(I)/nsp7 interface during nucleotide addition cycle. In the pre-translocated RdRp/RNA/Remdesivir complex, the nsp8(I)/nsp7 interface also exhibits similar movement, although to a lesser extent, compared to the apo RdRp [15], suggesting this movement might be necessary during multiple steps of the polymerization cycle.

Based on these structural and biochemical data, it is likely that a moderate interface between nsp8(I) and nsp7 may facilitate the formation of a relatively stable polymerase core and concurrently allow for certain flexibility of the nsp8(I)/nsp7 interface, which are critical for highly efficient and processive RNA synthesis. This nsp8(I)/nsp7 interface rearrangement appears to be translocation-coupled during nucleotide addition cycle and leads to a different interface between nsp8(I) and nsp7 subunits. Thus, we suggest that the nsp8(I)/nsp7 interface is an important regulatory site for RNA polymerase activity of RdRp, and this regulation mechanism might also be applicable to other coronaviruses due to the conservation in sequence and structure. In addition, small-molecule antiviral agents targeting this regulatory site of the SARS-CoV-2 RdRp complex hold advantages in specificity and efficacy over the conventional nucleoside inhibitors, whose anti-coronavirus potencies are undermined by the 3′ to 5′ proofreading exoribonuclease nsp14 [31]. Therefore, our findings elucidate new aspects of the mechanisms governing SARS-CoV-2 genome replication and suggest an alternative approach to target the viral polymerase by small-molecule drugs.

## MATERIALS AND METHODS

### Protein expression and purification

The full-length genes of SARS-CoV-2 nsp7 (residues 1-83), nsp8 (residues 1-198) and nsp12 (residues 1-932) (GenBank accession number MN908947) were all chemically synthesized with codon optimization (S2 Table) (GENEWIZ). The −1 ribosomal frameshifting that naturally occurs in virus to produce nsp12 was corrected in the synthesized gene to express nsp12 protein. The genes were cloned (S3 Table) into pET21a vector containing a C-terminal 6×His-tag. Mutations in nsp7 and nsp8 coding region in pET21a constructs were introduced following the Quickchange site-directed mutagenesis protocol (Stratagene). All constructs were transformed into *Escherichia coli* BL21(DE3) competent cells. The cells were grown in LB medium with 100 µg/mL ampicillin at 37 °C. After the OD_600_ reached 0.6, the culture was supplemented with 0.5 mM isopropyl-β-D-thiogalactopyranoside (IPTG) to induce protein expression at 16 °C for 16-18 h. The bacteria were then harvested by centrifugation, resuspended in lysis buffer containing 50 mM Tris-HCl, pH 8.0, 300 mM NaCl, 10 mM imidazole, 5% (v/v) glycerol, lysed by ultrasonication and then centrifuged at 80,000 g for 1 h using a T29-8×50 fixed angle rotor (Sorvall LYNX 6000 superspeed centrifuge, Thermo Scientific). The recombinant proteins were purified through 5 mL HisTrap HP column (GE Healthcare), and the eluted protein was further purified through 5 mL HiTrap Q HP column (GE Healthcare). Finally, the protein was applied to a gel filtration column, 120 mL HiLoad 16/600 Superdex 200 pg (GE Healthcare) in a buffer containing 20 mM Tris pH 7.5, 250 mM NaCl, 4 mM MgCl_2_. Purified proteins were concentrated to about 5 mg/ml, and stored at −80 °C until use.

### Assembly of RdRp/RNA complex

For assembling the nsp12-nsp7-nsp8/RNA complex, 5 µM purified nsp12 was incubated with nsp7, nsp8 and self-annealing RNA (RNA1, S3 Table) at a molar ratio of 1:2.5:2.5:3 at 30 °C for 10 minutes and then 4 °C overnight, in a buffer containing 20 mM Tris-HCl, pH 7.5, 250 mM NaCl and 4 mM MgCl_2_, 4 mM Dithiothreitol (DTT) and 100 µM ATP.

### Cryo-EM grid preparation and data acquisition

In total, 3.5 µl of the assembled complex was applied on Quantifoil R2/2 200 mesh Cu grids (EM Sciences) glow-discharged at 15 mA for 60 sec. The grid was then blotted for 3 sec at 4 °C under the condition of 100% chamber humidity, and plunge-frozen in liquid ethane using a Vitrobot mark IV (FEI). The grids were imaged using a 300 keV Titan Krios microscope (FEI) equipped with a Falcon III direct electron detector (FEI) at The Hormel Institute, University of Minnesota. Data were collected in counted mode with a pixel size of 0.89 Å and a defocus range from −1.0 to −2.4 µm using EPU (FEI). Each micrograph consists of 30 dose-framed fractions and was recorded with a dose rate of 0.8 e^-^/pixel/sec (1 e^-^/Å^2^/sec). Each fraction was exposed for 1 sec, resulting in a total exposure time of 30 sec and the total dose of 30 e^-^/Å^2^.

### Image processing

Cryo-EM data were processed using cryoSPARC v2.15 [32], and the procedure is outlined in S1A Fig. A total of 4,750 movies were collected. Beam-induced motion and physical drift were corrected followed by dose-weighting using the Patch motion correction [33]. The contrast transfer functions (CTFs) of the summed micrographs were determined using Patch CTF estimation [34]. From the summed images, particles were then automatically picked using Blob picker with the parameters: minimum particle diameter (60 Å) and maximum particle diameter (120 Å). In total, 4,052,050 particles were picked with a 216 pixels of box size. Three rounds of 2D classifications were performed to remove junk particles. Particles in good 2D classes (S1B Fig) were selected to generate multiple initial models using Ab-initio Reconstruction with five initial models for one calculation job and three initial ones for another job. The 20 Å low pass filtered initial models were set as the starting references for heterogeneous refinement (3D classification) in cryoSPARC v2.15. Particles in the best 3D classes from the two jobs were selected to perform homogeneous refinement, respectively. Particles from two homogeneous refinements were combined with duplicate particles removed using the Particle Sets Tool in cryoSPARC v2.15, and were subjected to a final non-uniform refinement in cryoSPARC v2.15. Fourier shell correlation (FSC) at the 0.143 cutoff resulted in an overall 3.4 Å resolution for the map (S1C Fig) [35]. Local resolution variation was estimated from the two half-maps in cryoSPARC v2.15 (S1D Fig).

### Model building and refinement

The initial model was generated by docking the individual domains of the apo RdRp of SARS-CoV-2 (PDB ID 6M71) into the cryo-EM density map using Chimera [36] and COOT [37]. The 3.4 Å cryo-EM density map allowed us to individually dock nsp12, nsp7 and nsp8 and manually build RNA duplex, followed by iterative cycles of manual adjustment using COOT. The RNA building was performed using the map at a lower contour than that of protein portion due to a lower occupancy of RNA substrate. Only one base overhang is present on the template RNA strand, therefore we built one nucleotide (ATP) incorporated to the 3′ end of primer on the basis that the assembled RdRp complex is active for *in vitro* analysis. The two base overhang on the primer RNA strand was not built due to poor density. The intact model was then refined using real_space_refine in Phenix [38]. In the real-space refinement, minimization global, local grid search, and adp were performed with the secondary structure, rotamer, and Ramachandran restrains applied throughout the entire refinement. The final model has good stereochemistry as evaluated by MolProbity [39]. 3DFSC calculation [40] shows the cryo-EM map has a sphericity value of 0.715, suggesting the map did sample majority of the angular space despite the minor preferred orientation problem (S1E Fig). The split cryo-EM maps were generated using color zone with 1.5 Å coloring radius in volume viewer of Chimera [36]. Map reconstruction quality was also evaluated by Mtriage [41] in Phenix (S1F Fig). The statistics of cryo-EM data collection, 3D reconstruction and model refinement were shown in S1 Table. All figures were created using ChimeraX [42].

### RdRp enzymatic activity assay

To anneal the RNA duplex, RNA primer and template were mixed at equimolar ratios (2 µM) in annealing buffer (10 mM Tris-HCl pH 8.0, 50 mM NaCl), which was denatured at 95 °C for 5 min and then cooled slowly to 16 °C. The purified nsp12 (1 µM) was mixed with nsp7 and nsp8 proteins at the ratio of 1:1:2 to form RdRp complex, which was incubated with 0.2 µM RNA duplex in Reaction buffer containing 20 mM Tris pH 8.0, 10 mM KCl, 6 mM MgCl_2_, 1 mM DTT at 30 °C for 5 min. To initiate RNA extension, 200 µM ATP was added into the reaction using a 5′ U_10_ RNA duplex as template and NTP containing 200 µM ATP, 200 µM UTP and 1 µCi of [α-^32^P]GTP was added into reaction using short RNA duplex. After incubation at 30 °C for 15 s, 30 s, 1 min, 2 min, 5 min, and 10 min, equal volume of Stop buffer (95% formamide, 30 mM EDTA) was added. RNA products were heated at 70 °C for 5 min and then analyzed on denaturing 20% Urea-PAGE. The FAM-labeled products were imaged via Amersham Typhoon scanner (GE Healthcare) and the radioactive labeled products were visualized by phosphorimaging.

### Gel mobility shift assay

For gel shift assay, RdRp complex was assembled as described in enzymatic activity analysis. After that, RdRp complexes at concentrations of 200 nM, 400 nM and 800 nM were incubated with 100 nM of FAM-labeled RNA duplex at 37 °C for 15 min. Samples were resolved using 6% native gel running in 0.5×TBE buffer at 100 V for 1 h in ice-bath. Then gels were scanned by Amersham Typhoon scanner (GE Healthcare).

### Data availability

The cryo-EM density map of our SARS-CoV-2 RdRp/RNA complex has been deposited in the Electron Microscopy Data Bank under the accession number EMD-22288. The corresponding atomic coordinates for the atomic model have been deposited in the Protein Data Bank under the accession number 6XQB.

## ACKNOWLEDGEMENTS

We thank the staff at the cryo-EM facility and instrument core facility in the Hormel Institute, University of Minnesota, for providing help. We also thank the Core Facility and Technical Support of Wuhan Institute of Virology for help in radioactive and fluorescent tests. B.L. and Y.H. initiated and designed the experiments. This work was supported by the COVID-19 Rapid Response grant from University of Minnesota and the starting-up funding from The Hormel Institute, University of Minnesota granted to B.L. and the Youth Innovation Promotion Association CAS (Y201750) to Y.H. W.S. performed protein sample preparations and assembly of the complexes used in the structure determination. W.S. and B.L. performed cryo-EM grid preparation, screening, and optimization. B.L. conducted high throughput data collection on Titan Krios. Yang Y., W.S. and B.L. performed image processing. W.S. and B.L. carried out model building and refinement. M.C. and W.Z. constructed mutations and purified proteins. M.C., W.Z. and Y.H. performed *in vitro* biochemical tests. W.S., S.C., Yi Y., Yang Y., Y.H. and B.L. analyzed data. W.S., Yang Y., Y.H. and B.L. wrote the manuscript with the contributions from all the authors.

## Supplemental Information for

**S1 Fig.**
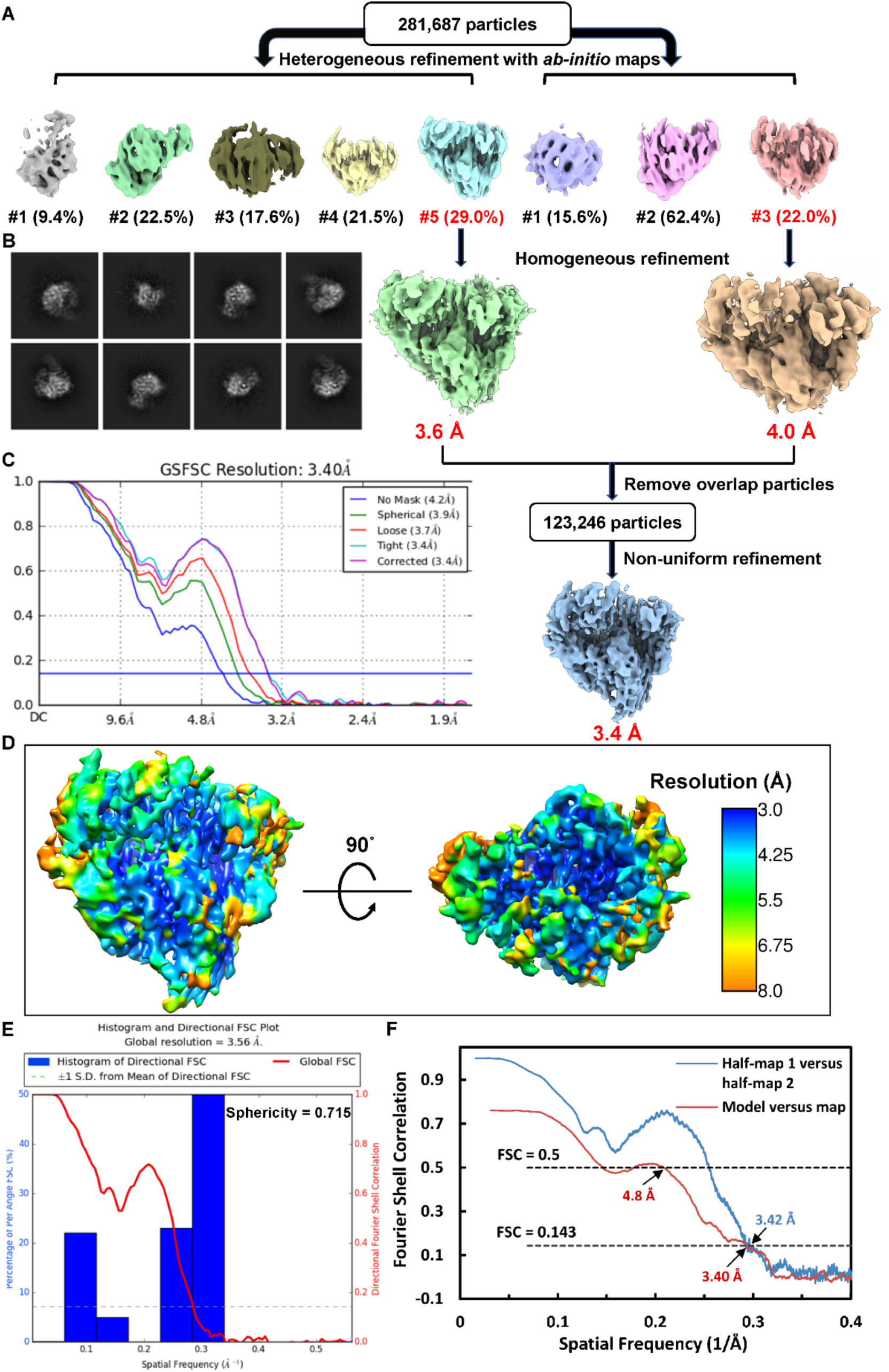
Cryo-EM data processing procedure for RdRp/RNA complex. **(A)** Flow chart of data processing and 3D reconstruction for the RdRp/RNA complex in the presence of ATP (see Methods details). **(B)** Representative 2D classes for initial model reconstruction. **(C)** Gold-standard Fourier Shell Correlations (FSCs) of the final map generated by cryoSPARC. **(D)** The overall local resolution map of the final 3D reconstruction. **(E)** Histogram and directional FSC plots of half maps with the corresponding sphericity value for the final 3D reconstruction. **(F)** FSC curves of half maps and model-to-map validated by Mtriage for the final 3D reconstruction.

**S2 Fig.**
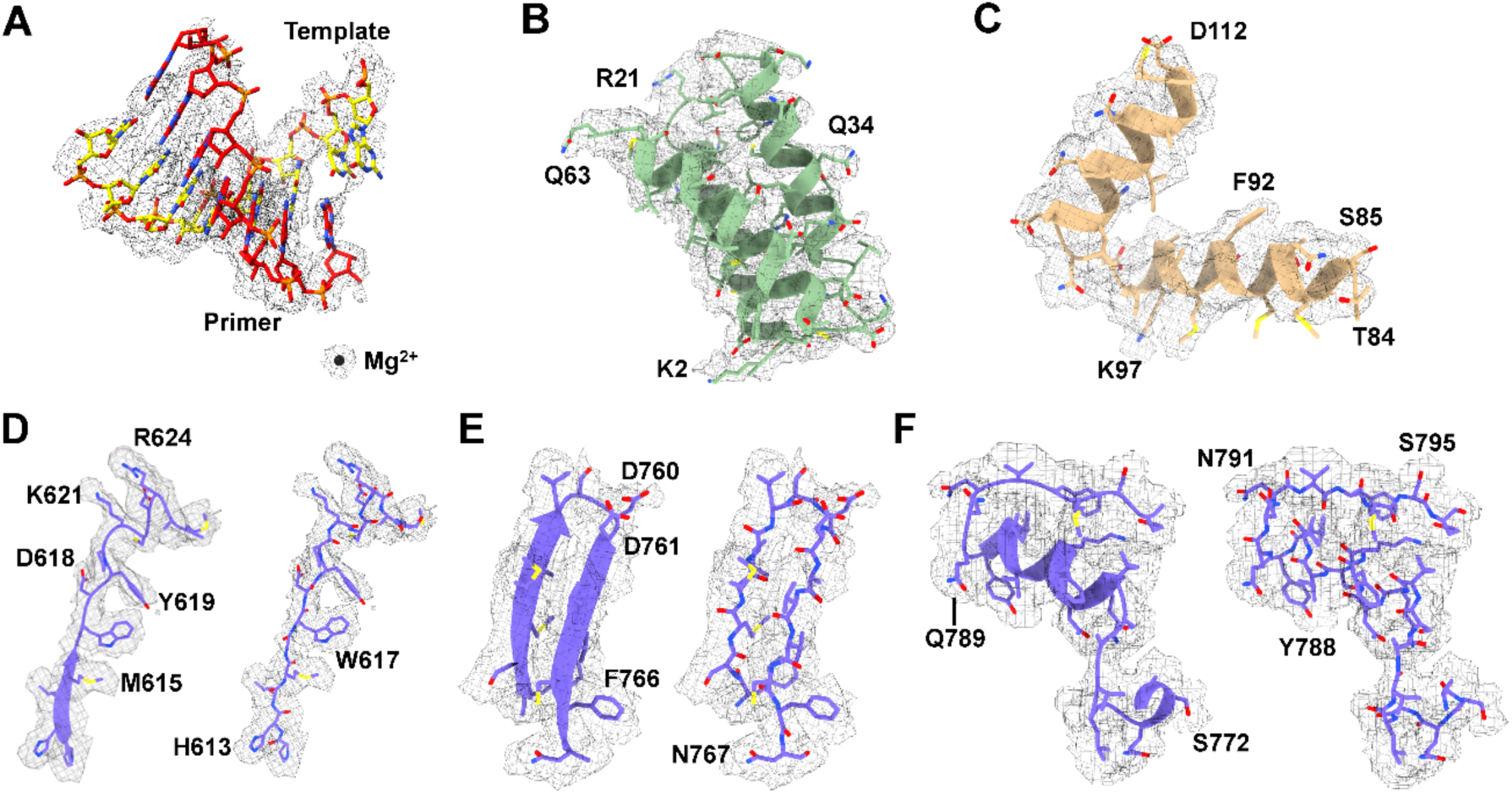
Cryo-EM maps for representative segments of RdRp/RNA complex. **(A-C)** Representative regions of the cryo-EM maps for duplex RNA (**A**), nsp7 (**B**) and nsp8(II) (**C**) cofactors in the complex. **(D-F)** Representative regions of the cryo-EM maps for motif A (**D**), motif C (**E**) and motif D (**F**) of nsp12 in the complex. The split cryo-EM density maps (mesh) fitted with atomic models (sticks) and contoured at 0.1 of view value of Chimera except for 0.08 used for the duplex RNA, demonstrating the agreement between the observed and modeled amino acid side chains.

**S3 Fig.**
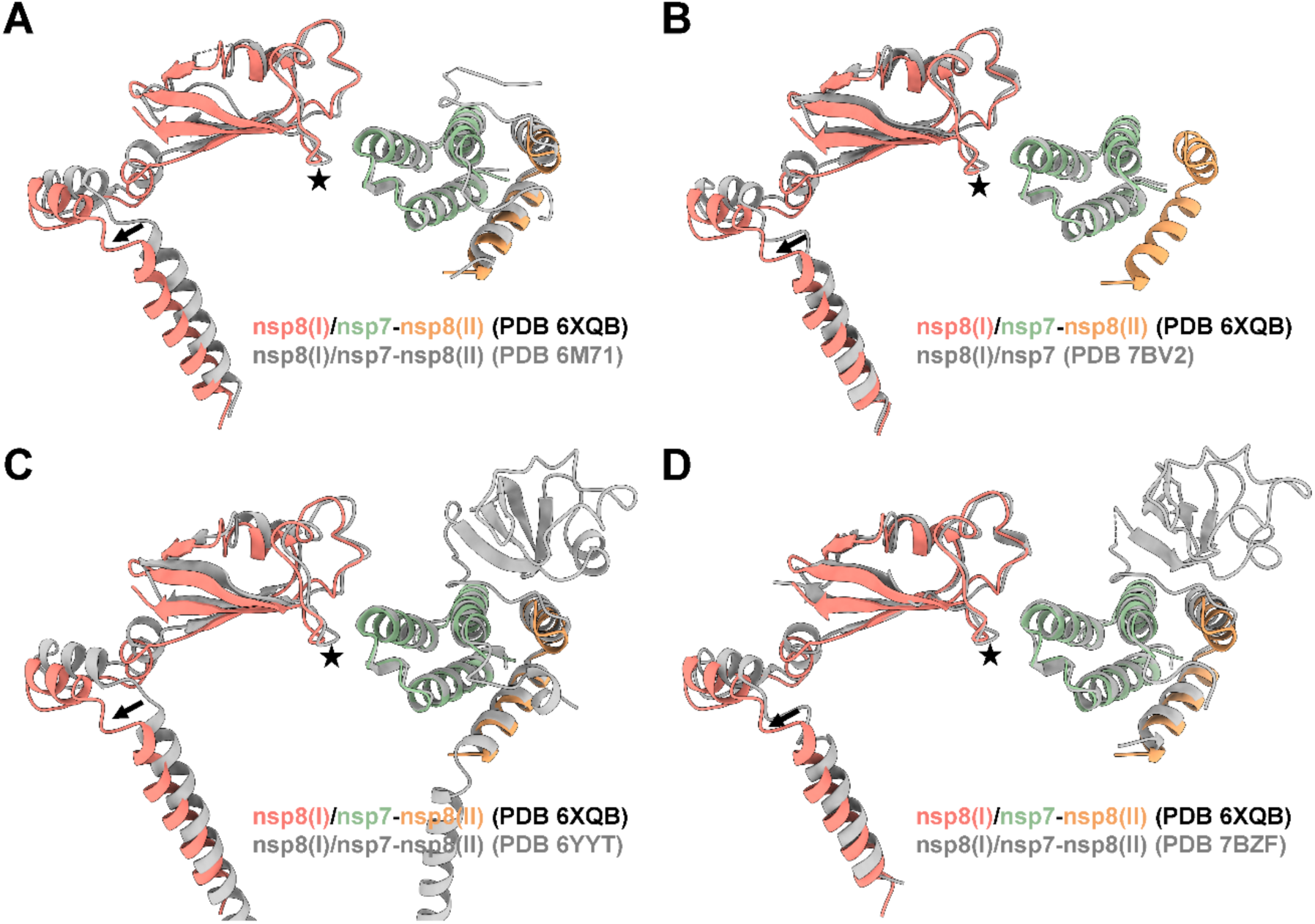
Superimpositions of the nsp8(I)/nsp7 interface among SARS-CoV-2 RdRp complexes. **(A-D)** Structural comparisons of the nsp8(I)/nsp7 interfaces among the different states of SARS-CoV-2 RdRp/RNA complexes: the post-translocated (PDB 6XQB, this study), the apo (PDB 6M71), the pre-translocated (PDB 7BV2), and other post-translocated (PDB 6YYT and 7BZF). The structures were superimposed on the nsp7. The nsp12 was omitted here for clear representation. The black stars mark the shift of nsp8(I) relative to nsp7 (∼1-2 Å).

**S4 Fig.**
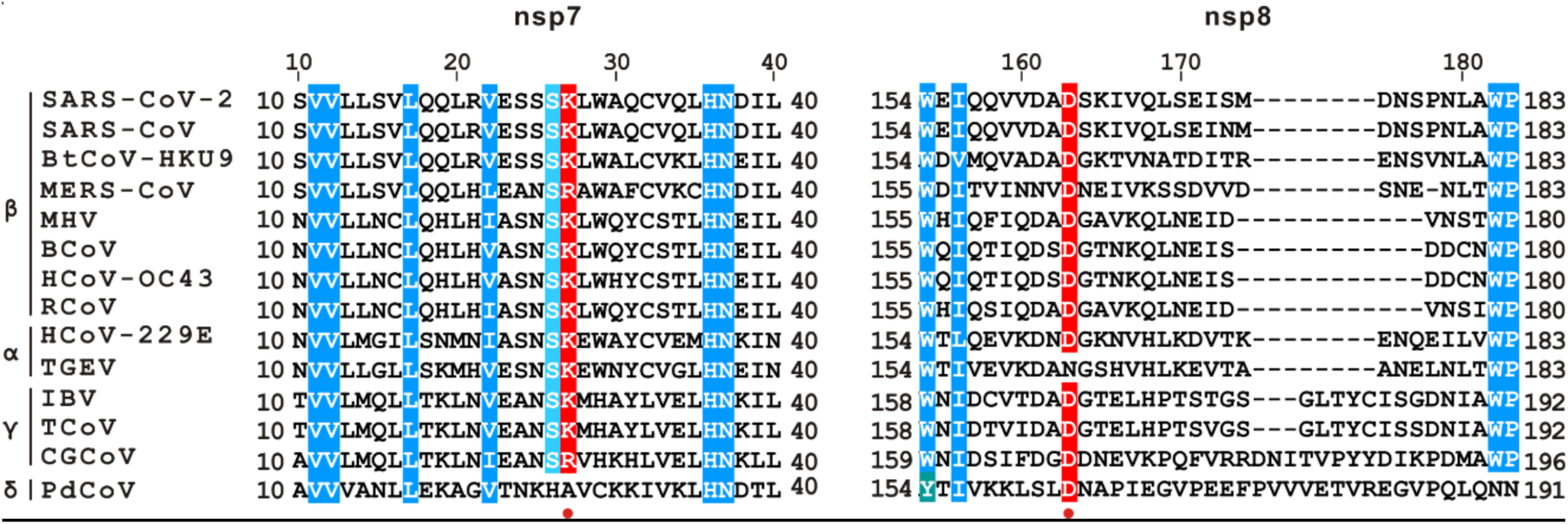
Sequence alignment of the nsp8(I)/nsp7 interface region among coronaviruses. Sequences of SARS-CoV-2 (MN908947), SARS-CoV (NP_828849), Bat coronavirus HKU9-1 (BtCoV-HKU9, ABN10910), Middle East respiratory syndrome-related coronavirus (MERS-CoV, QGV13494), Murine hepatitis virus (MHV, YP_009824978), Bovine coronavirus (BCoV, NP_150073), Human coronavirus OC43 strain (HCoV-OC43, YP_009555238), Rat coronavirus Parker (RCoV, YP_003029844), Human coronavirus 229E (HCoV-229E, AGW80667), Transmissible gastroenteritis virus (TGEV, ABC72413), Infectious bronchitis virus (IBV, YP_009824996), Turkey coronavirus (TCoV, YP_001941164), Canada goose coronavirus (CGCoV, YP_009755895), Porcine deltacoronavirus (PDCoV, QBQ34480) were used in ClustalW analysis. Accession codes in parentheses are for NCBI. Four genera of coronaviruses are indicated as α, β, γ and δ, respectively. The residues with direct contact in nsp8(I)/nsp7 interface are indicated by red dots.

**S5 Fig.**
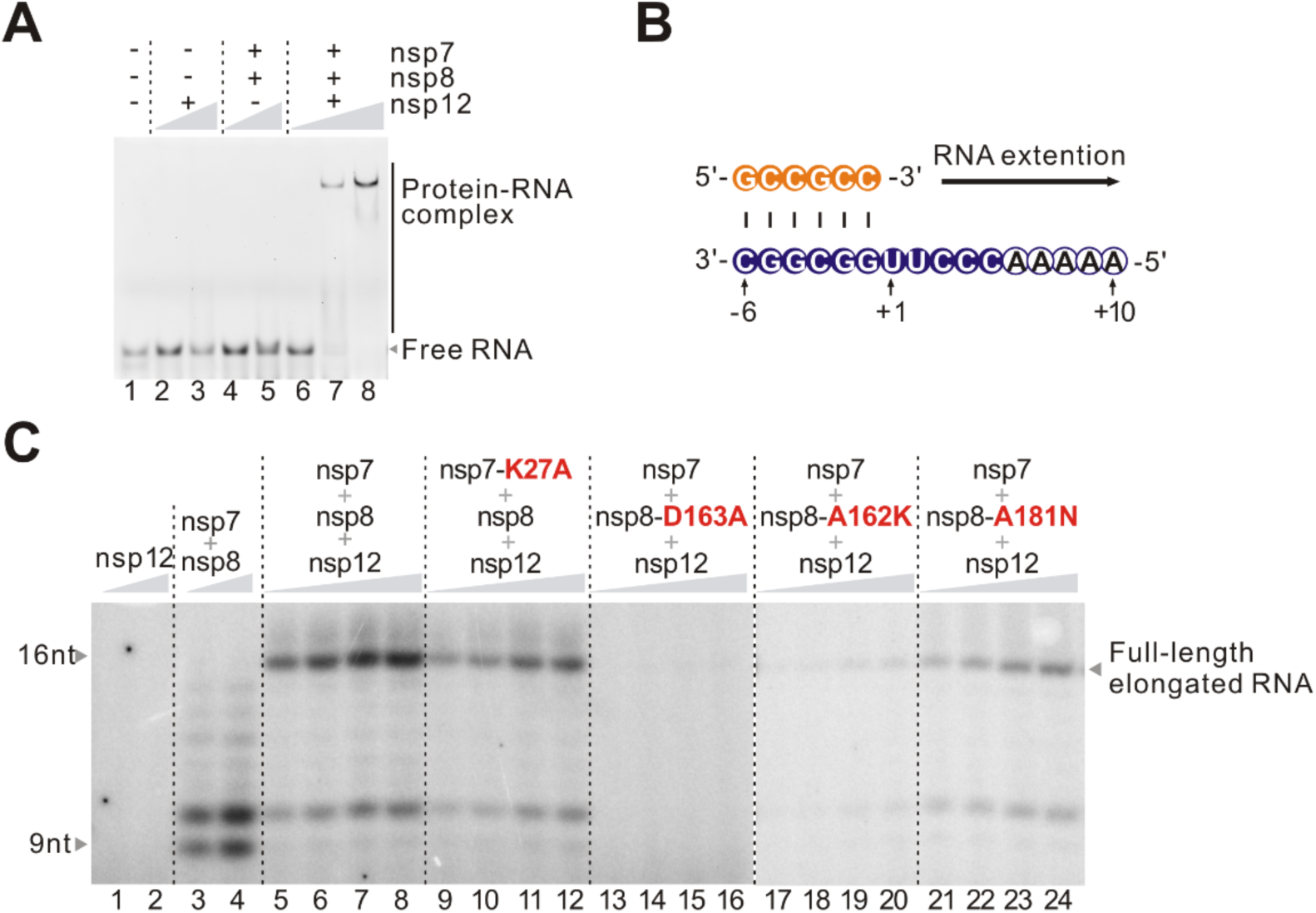
Contribution of nsp8(I)/nsp7 interface in RNA binding and extension of RdRp complex. **(A)** Gel mobility of RdRp complex and separated subunits in RNA binding. The nsp12 protein alone or nsp7-nsp8 complex were used at concentration of 400 nM and 800 nM. And the nsp12-nsp7-nsp8 complex was used at 200 nM, 400 nM and 800 nM. **(B)** Sequence of short RNA duplex used in RNA extension analyses. **(C)** Activity of recombinant RdRp complexes on short RNA hybrid shown in panel **B**. Reactions were carried out for 1, 2, 5 and 10 min at 30 °C. Reactions for nsp12 or nsp7-nsp8 complex were carried out for 5 and 10 min. Each test was repeated for three times and representative data are shown.

**S6 Fig.**
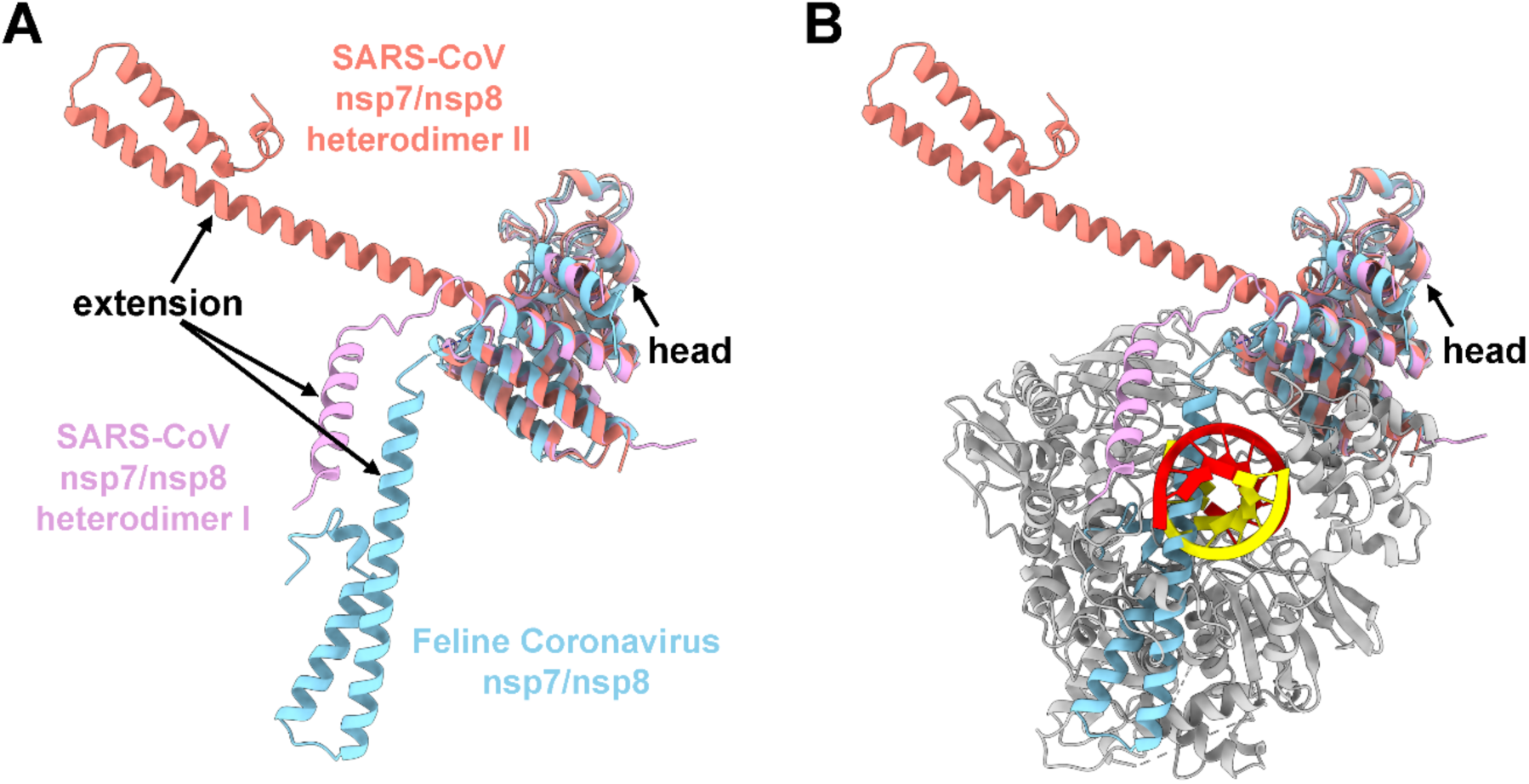
Superimposition of distinct nsp7-nsp8 heterodimers. **(A)** Superimposition of two heterodimers from SARS-CoV nsp7-nsp8 crystal structure (PDB 2AHM) and one heterodimer from Feline Coronavirus nsp7-nsp8 crystal structure (PDB 3UB0). Nsp7 and nsp8 head domain of three heterodimers superimpose well with each other, revealing a high flexibility of N-terminal extension in nsp8. **(B)** Superimposition of three heterodimer with the RdRp/RNA complex in this study. The flexibility of nsp8 N-terminal extension makes it readily interact with the upstream of RNA duplex.

**S1 Table.** Cryo-EM data collection, refinement and validation statistics.

| **Data collection/processing** | **SARS-CoV-2**<br>**nsp12-nsp7-nsp8/RNA complex** |
| --- | --- |
| Microscope | Krios |
| Voltage (kV) | 300 |
| Camera | Falcon III |
| Camera mode | Counting |
| Defocus range (μm) | −1.0 ~ −2.4 |
| Exposure time (s) | 30 |
| Dose rate (e<sup>−</sup>/pixel/s) | 0.8 |
| Magnified pixel size (Å) | 0.89 |
| **Reconstruction** |  |
| Software | cryoSPARC v2.15 |
| Symmetry | C1 |
| Particles refined | 123,246 |
| Resolution (Å) | 3.4 |
| Access code | EMD-22288 |
| **Model Statistics** |  |
| Number of residues (modeled) | 1,026 |
| Map CC (Chimera) | 0.7522 |
| MolProbity score | 2.00 |
| All-atom Clashscore | 6.13 |
| Cβ deviations | 0 |
| Rotamer outliers | 0.23% |
| Ramachandran |  |
| Outliers | 0.00% |
| Allowed | 15.15% |
| Favored | 84.85% |
| RMS deviations |  |
| Bond length | 0.009 |
| Bond angles | 1.161 |
| Access code | 6XQB |

**S2 Table.**
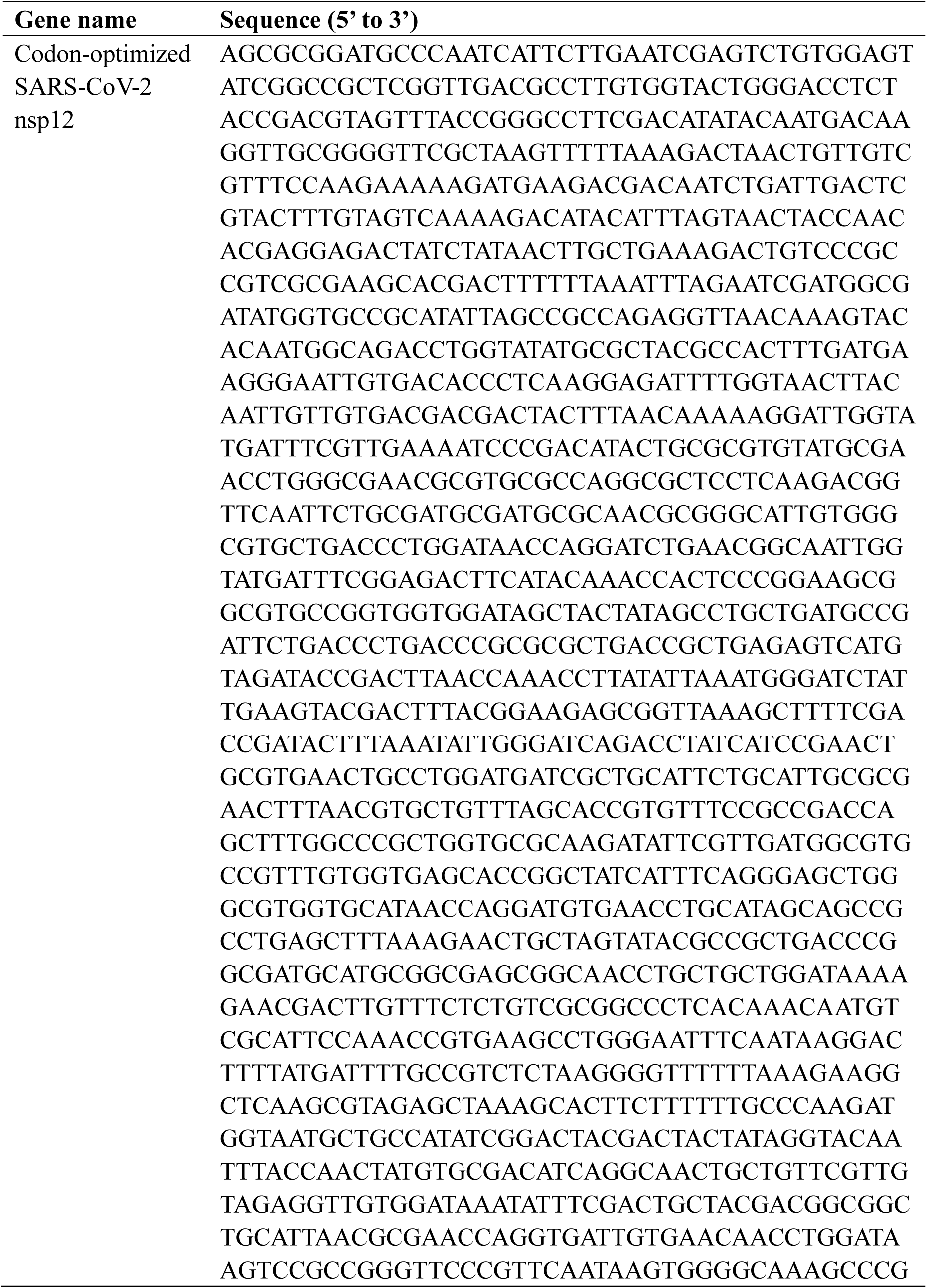
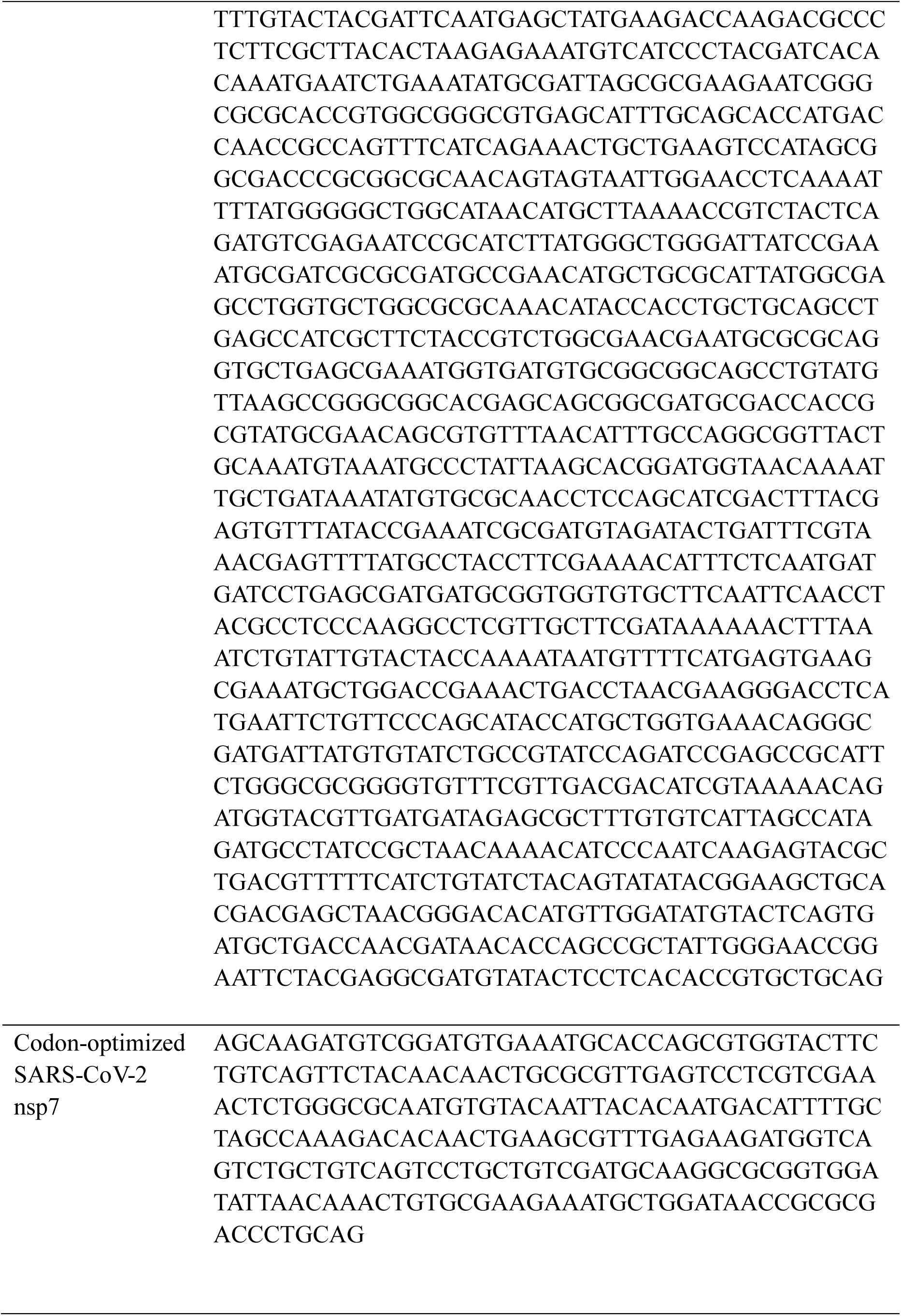
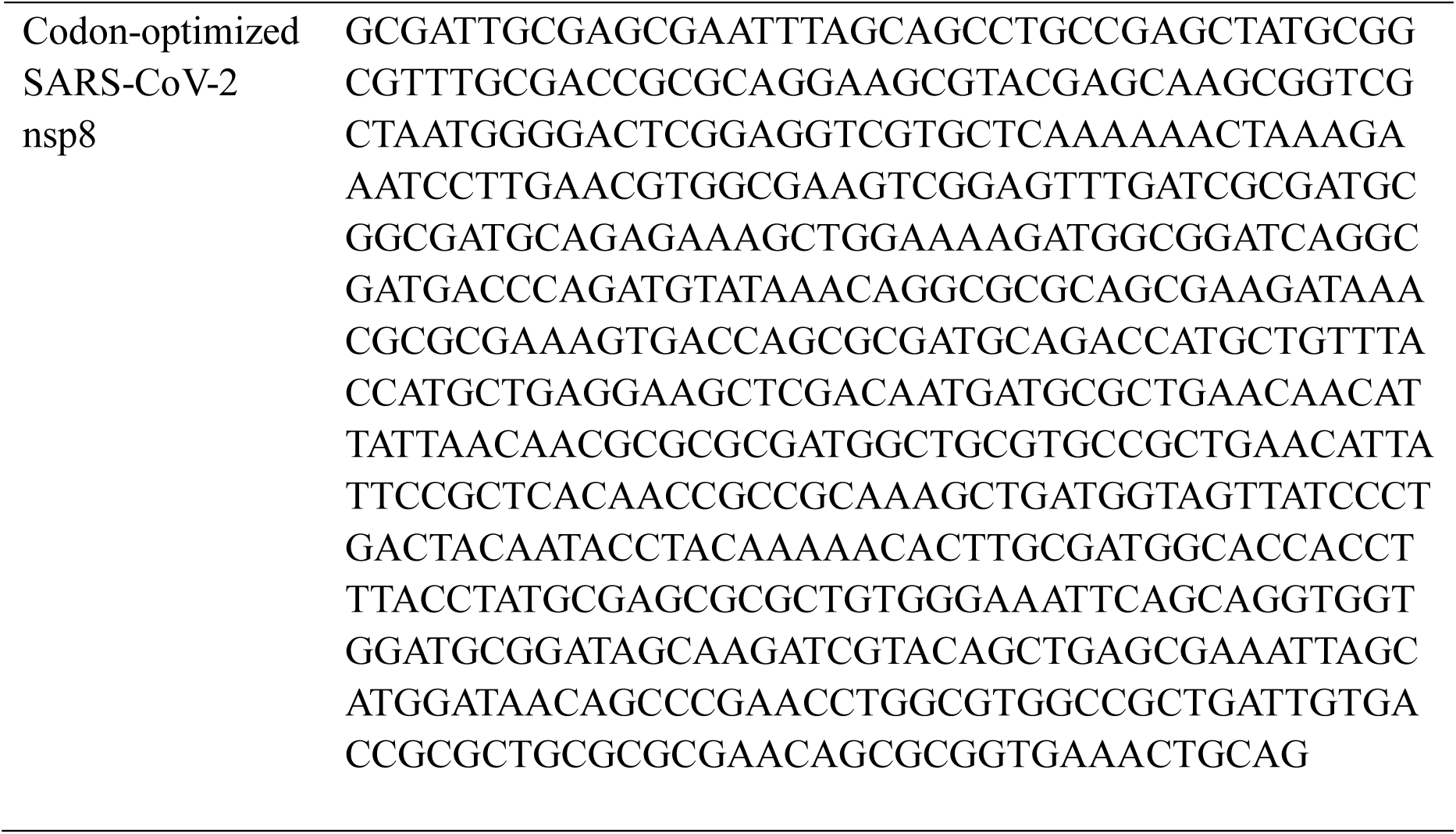
Codon-optimized nsp gene sequences used for expression.

**S3 Table.**
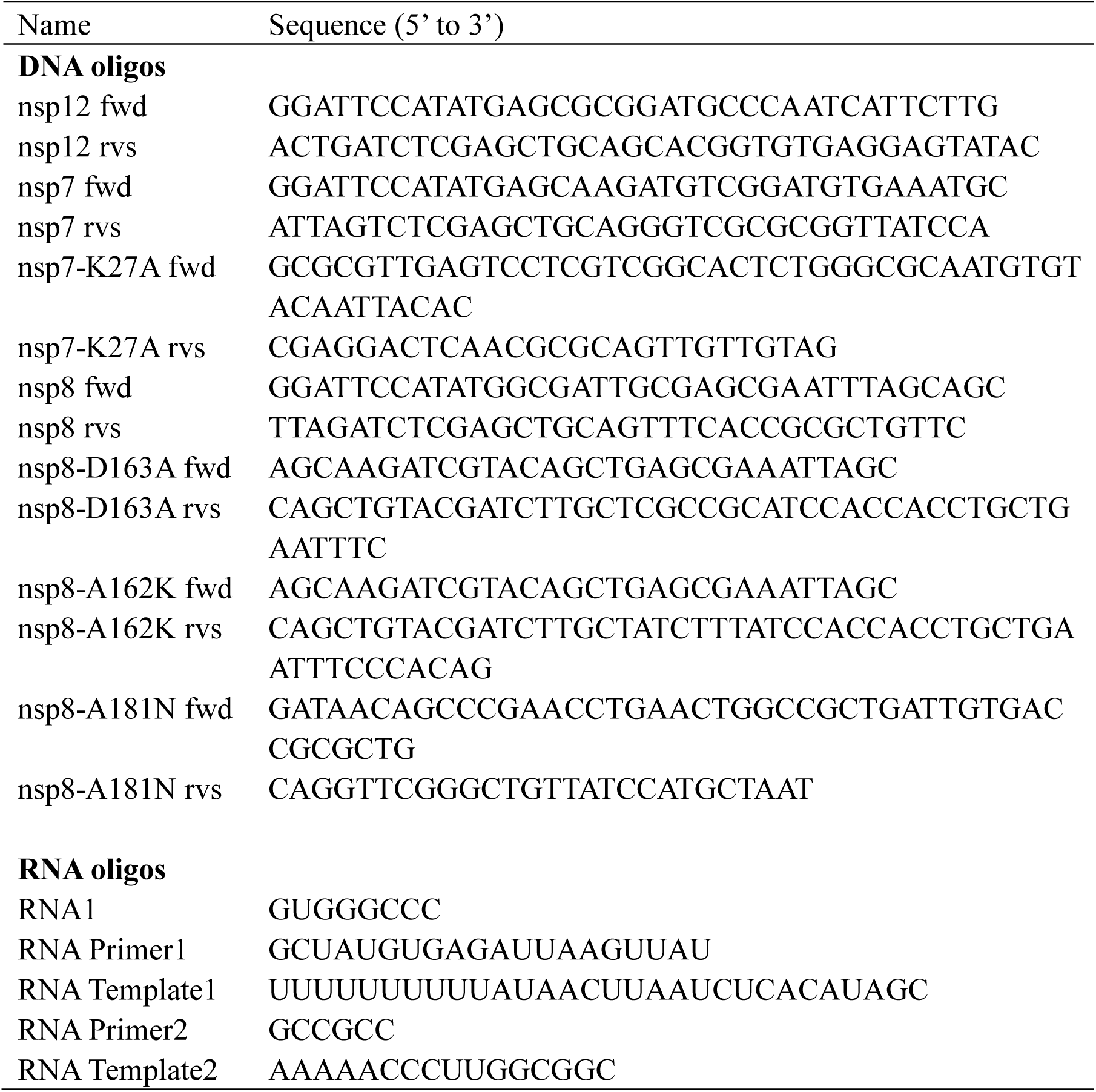
DNA primers used for cloning and RNA used in the study.

